# On the sensitivity analysis of porous finite element models for cerebral perfusion estimation

**DOI:** 10.1101/2021.02.18.431511

**Authors:** T. I. Józsa, R. M. Padmos, W. K. El-Bouri, A. G. Hoekstra, S. J. Payne

## Abstract

Computational physiological models are promising tools to enhance the design of clinical trials and to assist in decision making. Organ-scale haemodynamic models are gaining popularity to evaluate perfusion in a virtual environment both in healthy and diseased patients. Recently, the principles of verification, validation, and uncertainty quantification of such physiological models have been laid down to ensure safe applications of engineering software in the medical device industry. The present study sets out to establish guidelines for the usage of a three-dimensional steady state porous cerebral perfusion model of the human brain following principles detailed in the verification and validation (V&V 40) standard of the American Society of Mechanical Engineers. The model relies on the finite element method and has been developed specifically to estimate how brain perfusion is altered in ischaemic stroke patients before, during, and after treatments. Simulations are compared with exact analytical solutions and a thorough sensitivity analysis is presented covering every numerical and physiological model parameter.

The results suggest that such porous models can approximate blood pressure and perfusion distributions reliably even on a coarse grid with first order elements. On the other hand, higher order elements are essential to mitigate errors in volumetric blood flow rate estimation through cortical surface regions. Matching the volumetric flow rate corresponding to major cerebral arteries is identified as a validation milestone. It is found that inlet velocity boundary conditions are hard to obtain and that constant pressure inlet boundary conditions are feasible alternatives. A one-dimensional model is presented which can serve as a computationally inexpensive replacement of the three-dimensional brain model to ease parameter optimisation, sensitivity analyses and uncertainty quantification.

The findings of the present study can be generalised to organ-scale porous perfusion models. The results increase the applicability of computational tools regarding treatment development for stroke and other cerebrovascular conditions.

## 1 Introduction

Cerebrovascular diseases including stroke impose a heavy burden on society [1]. The majority of stroke cases are caused by a thrombus blocking a major cerebral artery leading to severe blood shortage in the brain (ischaemic stroke). In recent decades, ischaemic stroke treatment has been revolutionised by thrombolysis (dissolving of blood clot with a thrombolytic agent) [2] and thrombectomy (mechanical thrombus removal with a stent retriever) [3, 4]. Advancing the related clinical procedures has great potential to benefit stroke patients but is tied entirely to resource intensive animal experiments and clinical trials [3–7]. To ease this task, the *In Silico* clinical trials for the treatment of acute Ischaemic STroke (INSIST) consortium aims to develop computational models of stroke and its treatments [8]. Once reliable models are available, *in silico* clinical trials could sharpen the focus of clinical trials and hence save time and money [9–11].

Estimating brain perfusion in both healthy and occluded scenarios is an important element of the INSIST pipeline and serves as a bridge between localised treatment effects and patient outcome [8]. To this end, a multi-scale haemodynamic model is under development which combines a one-dimensional (1D) network model of large arteries [12, 13] and a three-dimensional (3D) porous perfusion model [14] to simulate blood flow in the entire vasculature. The models will be coupled strongly through the cortical surface. The envisaged interface will facilitate communication between the boundary conditions of the models, namely pressure and volumetric flow rate values averaged over cortical territories [13]. Previously somewhat similar weak-coupling schemes have been established between a lumped parameter model and a porous continuum model to cover arteriole boundary conditions at the brain surface [15–19]. This approach preserves the two models as independent functional units and avoids the positioning of volumetric sources required for volumetric coupling [20–22]. The anatomical connection between large arteries and the microcirculation is crucial in ischaemic stroke modelling because it determines the location and extent of ischaemic regions. Another advantage of placing the interface at the cortical surface is that anatomical connections are preserved at the cortical surface so that subcortical vessels can be homogenised to lower computational costs. In the case of volumetric coupling, subcortical arterioles must be added to the one-dimensional network model to account for such connections which can drastically increase the number of discretised branches.

Computational porous and poroelastic models have been applied over a number of decades and have been proposed to describe aspects of cardiac [23–25] and cerebral biomechanics [15–19, 26]. Such organ-scale computational models have been applied to capture both healthy and pathophysiological states, such as brain injury [27–29], oedema [30], conditions associated with dementia [15, 18, 19], and ischaemic stroke [14]. The majority of these models utilise the Finite Element (FE) method [14–19, 27–30] with relatively few exceptions [20–22]. The present study focuses on a porous perfusion FE model developed specifically to capture blood flow changes in ischaemic stroke [14] as part of the *in silico* trial pipeline of INSIST [8]. Even though organ-scale porous and poroelastic brain models are becoming increasingly popular [14–19, 22, 26–32], efforts regarding their verification, validation [15], comprehensive sensitivity analysis [22, 33], and uncertainty quantification [31, 32] remain relatively rare and there is thus a need to fill this gap before the models are applied for clinical trial design and decision making.

Recently, the American Society of Mechanical Engineers (ASME) has introduced the Verification and Validation (V&V) 40 standard (titled “Assessing Credibility of Computational Modeling through Verification and Validation: Application to Medical Devices”) [34], which details the requirements that a credible engineering software needs to satisfy to ensure safe applicability in the medical device industry. The present study thus sets out to establish guidelines regarding the efficient usage of a porous cerebral perfusion Finite Element model [14] both in terms of accuracy and computational cost. To this end, simulations will be compared to exact analytical solutions and a thorough sensitivity analysis will be carried out covering every numerical and physiological model parameter. The results prepare the ground for uncertainty quantification, virtual patient generation, and validation, which are essential to carry out reliable *in silico* trials of ischaemic stroke treatments.

## 2 Methodology

Inspired by recent advances in the fields of heart [23–25] and brain [15–19] haemodynamic modelling, a multi-compartment porous continuum framework is utilised to describe time-averaged cerebral blood flow in the microcirculation embedded within brain tissue. In this approach, microscale individual blood vessels are not captured. Instead, porous media are introduced to describe characteristic units of the vessel network, such as bundles of penetrating arterioles and the capillary mesh as shown in Fig. 1. During the macroscale continuum description, the corresponding vessels are grouped into multiple compartments (Figs. 1b, c, and d). In practice, the corresponding vessels can be isolated, for instance, on a geometrical basis using diameter thresholding or branching order, or on a structural basis considering layers of the vessel walls (smooth muscle cells and endothelial cells). The remaining vessels, which are not part of any of these compartments and are not shown in Fig. 1, serve as a link between the compartments.

**Figure 1:**
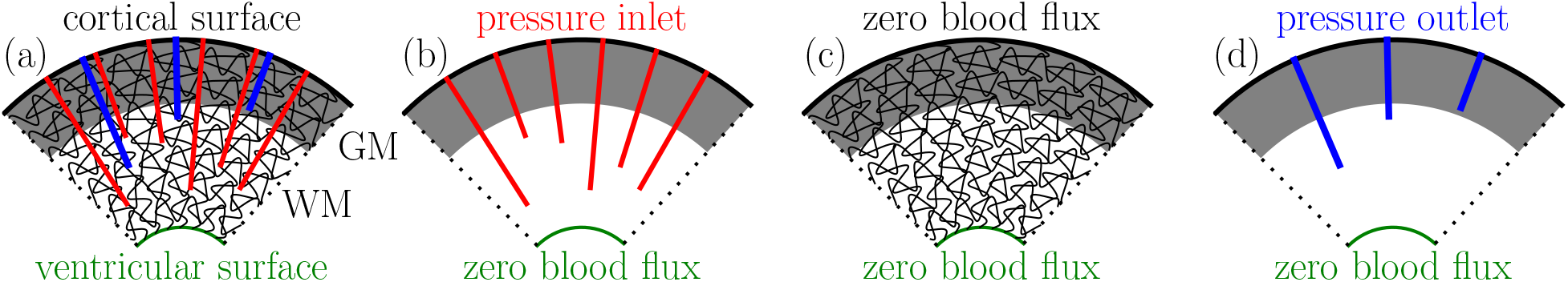
Schematic of the microcirculation along a slice in a cortical column: descending arterioles (red), capillaries (black) and ascending veins (blue) are shown in both grey (GM) and white matter (WM) (a). The porous cerebral blood flow model relies on the idea of separating these vessels into coexisting arteriole (b), capillary (c), and venule (d) compartments.

In the porous formulation, the primary flow variable is the volume-averaged (Darcy) pressure which can be deduced from the pressure distribution of the individual blood vessels by spatial averaging. The general partial differential equation set describing a steady-state multi-compartment porous Darcy system [15–19, 23–25] is written in a compact form^1^ as

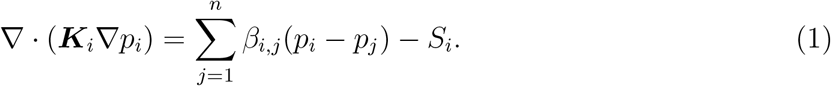

Here, *p_i_* denotes the Darcy pressure in the *i*^th^ compartment and ***K**_i_* is the corresponding permability symbolising flow conductance through pores (lumen of vessels) with well-separated length scales. Considering *n* compartments, *i* =1, 2, … *n* and *β_i,j_* is the intercompartment coupling coefficient matrix with *n* × *n* elements representing conductance between compartments *i* and *j*. It follows that *β_i,j_* is arbitrary if *i* = *j*. A volumetric source term in the *i*^th^ compartment is defined by *S_i_*.

Considering a domain of interest denoted by Ω with a boundary *∂*Ω = Γ = Γ*_D,i_* ∪ Γ*_N,i_*, a set of Dirichlet type boundary conditions (DBC) and Neumann type boundary conditions (NBC) are formulated as

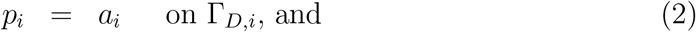

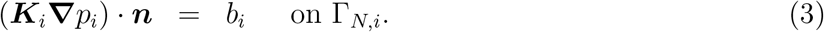

Here, ***n*** is the outward-pointing unit vector perpendicular to the boundary surface. DBCs can be used to describe, for example arterial blood pressure along the cortical surface as shown in Fig. 1a and b. By comparison, NBCs are typically used to enforce a specific volumetric blood flow rate across boundary regions, such as zero blood flux through the ventricular surface in Fig. 1b, c, and d. (NBC could be also used to account for cerebrospinal fluid filtration at the choroid plexus.)

Thereafter, the weak form can be derived by multiplying every component of Eq. (1) with a non-zero test function *t_i_*, integrating over Ω and applying the divergence theorem:

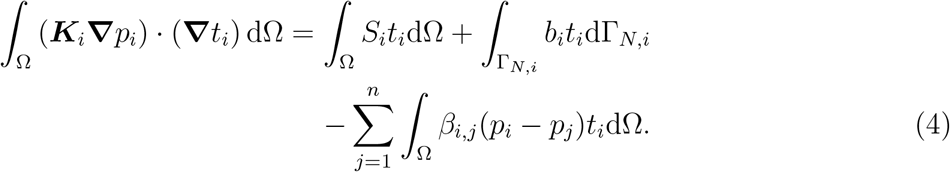

Eq. (4) is discretised by the continuous Bubnov-Galerkin method [35]. The *H*^1^ Sobolev space contains both the test and the trial functions so that

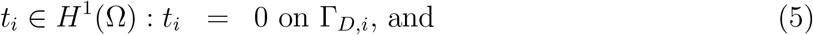

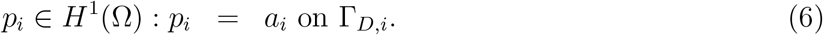

Numerical solutions of *p_i_* are obtained with the open source FEniCS library [36–38] using first or second order Lagrange elements denoted by *P*_1_ and *P*_2_, respectively. The permeability tensors ***K**_i_* and the coupling coefficients *β_i,j_* are captured by piecewise constant, zeroth order discontinuous elements (*dP*_0_) unless stated otherwise. For simplicity, we restrict ourselves to a model with arteriole (*i* = 1 = *a*), capillary (*i* = 2 = *c*), and venule (*i* = 3 = *v*) compartments, and then generalise our findings whenever it is possible (1). In this case, the remaining pre-capillary and post-capillary vessels are lumped into the arteriole-capillary (*β*_1,2_) and capillary-venule (*β*_2,3_) coupling coefficients, respectively.

Numerical solutions of linear equation systems originating from the FE discretisation are obtained iteratively based on the BiCojungate Gradient STABilised method (BiCGSTAB) [39]. Computations are accelerated with an Algebraic MultiGrid (AMG) preconditioner [40]. Simulations were run on a desktop with an Intel Xeon E-2146G processor and 32 GB RAM unless stated otherwise. In order to verify the resulting model and carry out a comprehensive sensitivity analysis, three test cases are considered as detailed in the following subsections.

### 2.1 Three-dimensional synthetic data

To derive manufactured solutions [41], we assume solution functions of Eq. (1) of the form

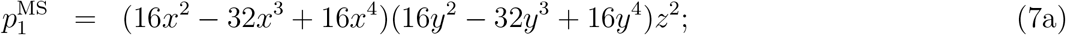

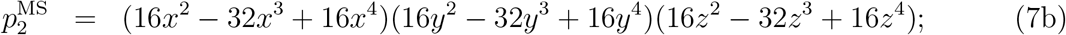

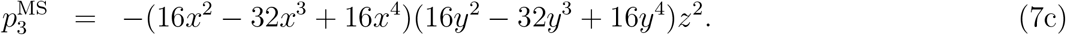

The domain of interest chosen here is a unit cube Ω = [0, 1] × [0, 1] × [0, 1] with periodic boundary conditions applied on four sides:

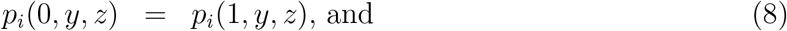

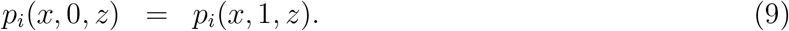

Eqs. (8) and (9) define four DBCs in every compartment and another one is added in the arteriole (*i* = 1) and the venule (*i* = 3) compartments based on the manufactured solutions such that

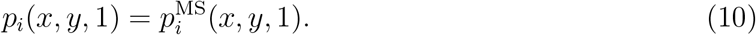

Thereafter, the equation set is closed by prescribing homogeneous NBCs on the remaining surfaces (in Eq. (3) *b_i_* = 0).

The permeability tensors and coupling coefficient matrices corresponding to the manufactured solutions are detailed in Appendix A. The chosen polynomial functions (7a)–(7c) are straightforward to differentiate and are the simplest functions which satisfy the imposed boundary conditions. The source terms *S*_1_, *S*_2_ and *S*_3_ satisfying Eq. (1) can be obtained by substituting Eqs. (7) and (31)–(32) into Eq. (1). The source terms are set to zero in every other test case (details in Sections 2.3 and 2.2).

### 2.2 Three-dimensional human brain

Details of the baseline human brain simulation are provided in [14] and therefore only the most important settings are covered here. The computational domain (Ω) is obtained by postprocessing a tetrahedral patient-specific head model utilised in multiple recent studies [14, 42–44]. The geometry depicted in Fig. 2a is remeshed with tetrahedral elements using Tetgen [45]. The volumetric region of interest includes both grey matter (Ω*_G_*) and white matter (Ω*_W_*) subdomains, so that Ω = Ω*_G_* ∪ Ω*_W_* as depicted in Fig. 2c. The bounding surface regions (∂Ω) include a transverse cut-plane of the brainstem Γ*_BS_*, the ventricles Γ*_V_* and the pial surface Γ*_P_* so that *∂*Ω = Γ*_BS_* ∪ Γ*_V_* ∪ Γ*_P_* as depicted in Figs. 2a and b. The boundary region associated with the pial surface is subdivided into eight perfusion territories corresponding to major feeding arteries as shown in Fig. 2a. Perfusion territories have been identified using a voxelised atlas created based on vessel-encoded arterial spin labelling (ASL) perfusion magnetic resonance imaging (MRI) [46–50]. Then, the surface region that is perfused, for instance, by the Right Middle Cerebral Artery (R-MCA) is denoted by Γ_R-MCA_. This approach ensures that blood arrives to the domain through cortical surface regions mirroring anatomical connections between large arteries and the microcirculation as shown in Fig. 2a.

**Figure 2:**
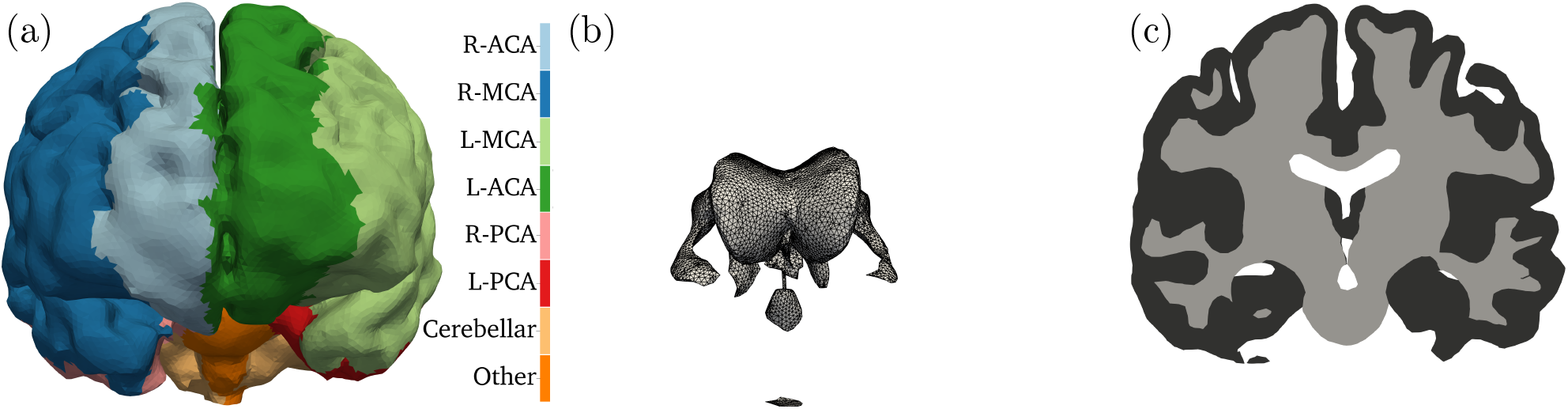
Coronal views of boundary regions and subdomains of the human brain model [44]: (a) pial surface; (b) ventricles and cut-plane at brainstem; (c) GM and WM along a slice.

The imposed baseline boundary conditions prescribe zero flow through the transverse cut-plane of the brainstem and the surface of the ventricles:

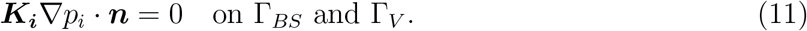

Flow through the pial surface in the capillary compartment is zero:

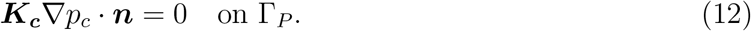

Because of the incompressible fluid flow model, the results depend solely on the cerebral perfusion pressure (CPP) defined as the arteriole-venule pressure difference at the pial surface: CPP = *p_a_* – *p_v_*. Setting the zero level of the pressure at the outlet of the venous compartment (*p_v_*) leads to

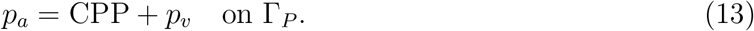

Pressure values in the case of 3D brain simulations are presented relative to the venous outlet pressure *p_v_*. To account for totally occluded scenarios, blood flow through the perfusion territory of an occluded vessel is set to zero whereas surface pressure is assumed to remain constant in other regions. Accordingly, a R-MCA occlusion is modelled with zero flux through the corresponding cortical territory (Γ_R-MCA_) so that the resulting mixed boundary conditions become

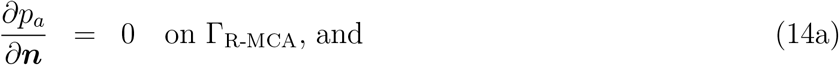

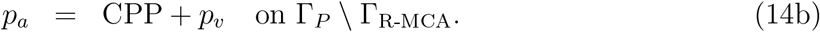

The model is parametrised based on several simplifications as discussed in [14]. Considering that the arteriole-capillary 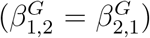 and capillary-venule 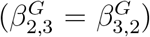 coupling coefficients are known in the grey matter, it is assumed that the ratio of grey and white matter coupling coefficients (*β^G^*/*β^W^*) is constant. It has been demonstrated based on microscale one-dimensional haemodynamic network simulations that the capillary permeability tensor is isotropic and characterised by a scalar *k*_2_ [51]. It is hypothesised that a reference coordinate system [*ξ*, *η*, *ζ*] can be found at every point so that *ζ* is parallel to the local axis of the descending arterioles and ascending venules. In this coordinate system, the anisotropic arteriole 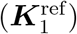 and venule 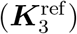 permeability tensors are modelled as tensors with a single non-zero diagonal element (*k*_1_ and *k*_3_) because they represent vessel bundles:

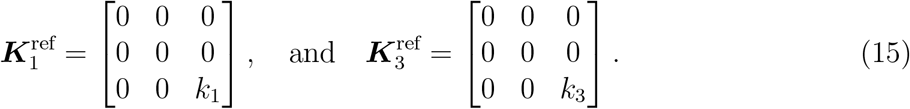

The [*ξ*, *η*, *ζ*] coordinate system is determined assuming that penetrating vessels grow from the pial surface to the ventricles following a vector field. This vector field is computed as a gradient of a scalar field governed by a single diffusion equation [14]. It is worth noting that no other methods have been proposed to obtain these anisotropic permeability fields even though they play a crucial role in predicting perfusion response to vessel occlusion.

### 2.3 One-dimensional brain tissue column

Eq. (1) can be simplified so that a one-dimensional problem with homogeneous permeabilities can be posed and defined by a set of ordinary differential equations in the form of

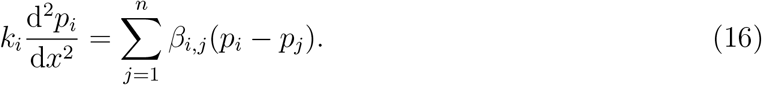

Here, the permeability of each compartment (*k_i_*) is a scalar. Introducing the dimensionless variables *x\** = *x/l_s_* and *p*\* = *p/p_s_* based on a length scale (*l_s_*) and a pressure scale (*p_s_*) leads to a dimensionless form of Eq. (16) as

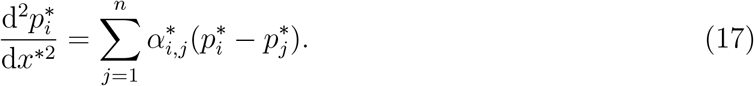

The dimensionless matrix group 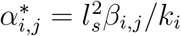 provides a similarity condition for Eq. (16).

Equation (17) describes *n* second order ordinary differential equations. This system can be replaced by 2*n* first order ordinary differential equations once the pressure gradient 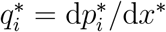 is introduced. Thereafter, the unknown vector

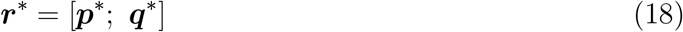

contains the pressure 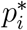 and pressure gradient 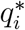 functions for each compartment. Finally, a matrix differential equation based on Equation (17) can be formulated as

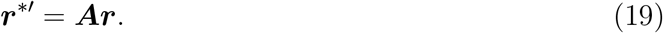

Here, the prime (′) denotes spatial differentiation and the ***A*** matrix is

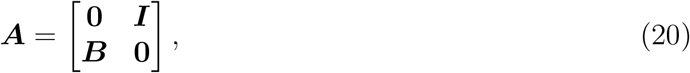

where every submatrix has a dimension of *n* × *n*, and ***I*** is the identity matrix. The connection between the coupling coefficient matrix ***α*** = *α_i,j_* and the submatrix ***B*** is given by

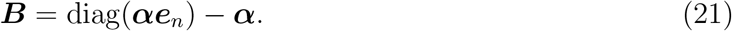

Here, the *n* dimensional vector ***e**_n_* is filled with ones, and the “diag” operator is used to turn a column vector into a diagonal square matrix.

The solution of Equation (19) is

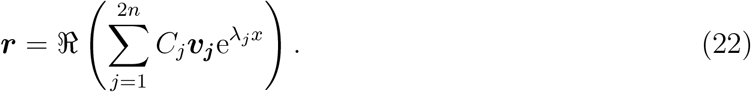

Here, λ*_j_* and ***v_j_*** are the *j*^th^ eigenvalue and eigenvector of ***A***, respectively. 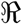 is used to take the real part of the expression and to handle complex eigenvalue and eigenvector pairs.

The *C_j_* coefficients are determined based on combinations of DBCs and NBCs at *x_b_*. A DBC in compartment *i* reads as

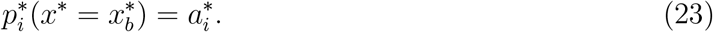

Based on the unknown vector defined in Eq. (18) and the solution expression Eq. (22), a DBC can be satisfied by imposing

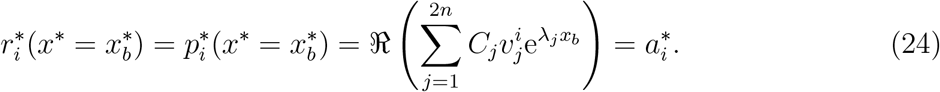

Here, 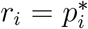 denotes the *i*^th^ element of ***r*** and 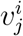 is the *i*^th^ element of *j*^th^ eigenvector. A NBC in compartment *i* is defined as

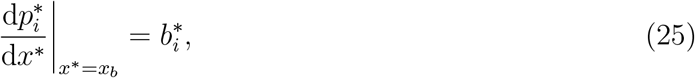

The second half of the unknown vector defined in Eq. (18) includes the pressure gradients. Therefore, a NBC can be satisfied by imposing

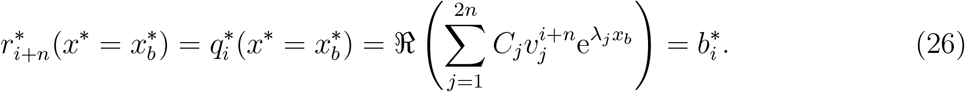

Eqs. (24) and (26) provide a single equation for each boundary condition. Considering *n* compartments, the 2*n* boundary conditions lead to 2*n* linear equations based on the 2*n* coefficients of the solution expressions (*C_j_*). The resulting linear equation system determines uniquely the coefficients of the solution expressions (*C_j_*). Using the DBC (23) and the NBC (25), the procedure can be extended to handle multiple subdomains with different coupling coefficients. In such cases, interface conditions have to be applied to ensures that the solution is continuously differentiable. Therefore, this analytical method is suitable to obtain solutions for a column of brain tissue stretching between the cortical and ventricular surfaces. The method can handle a domain including grey and white matter with identical permeabilities but different coupling coefficient matrices.

## 3 Results and Discussion

### 3.1 Reflection on the ASME V&V40 framework

The INSIST software suite [8] aims to quantify the efficacy of thrombolysis [2] and thrombectomy [3, 4] using computational models. Following the framework of the ASME V&V40 standard shown in Fig. 3, the **question of interest** can be phrased as “what is the best available treatment option in terms of functional outcome using stent retrievers and tissue plasminogen activator?” The **context of use** of the porous finite element model is quantifying how cerebral blood flow is altered in the microcirculation during ischaemic stroke and post-treatment compared to the healthy baseline case. The output of the FE model is the spatial distribution of brain tissue perfusion, which can be measured, for example, by ASL perfusion MRI in clinical settings [47]. The porous FE model provides (i) pressure and volumetric blood flow rate inputs for haemodynamic simulations in large arteries [13] which evaluate forces on the thrombus; and (ii) perfusion input for tissue health computation used for infarct volume estimation [52]. Therefore, the output of the FE model is linked indirectly to a statistical module estimating the functional outcome of individual virtual patients based on the computed infarct volume, and other features, such as age.

**Figure 3:**
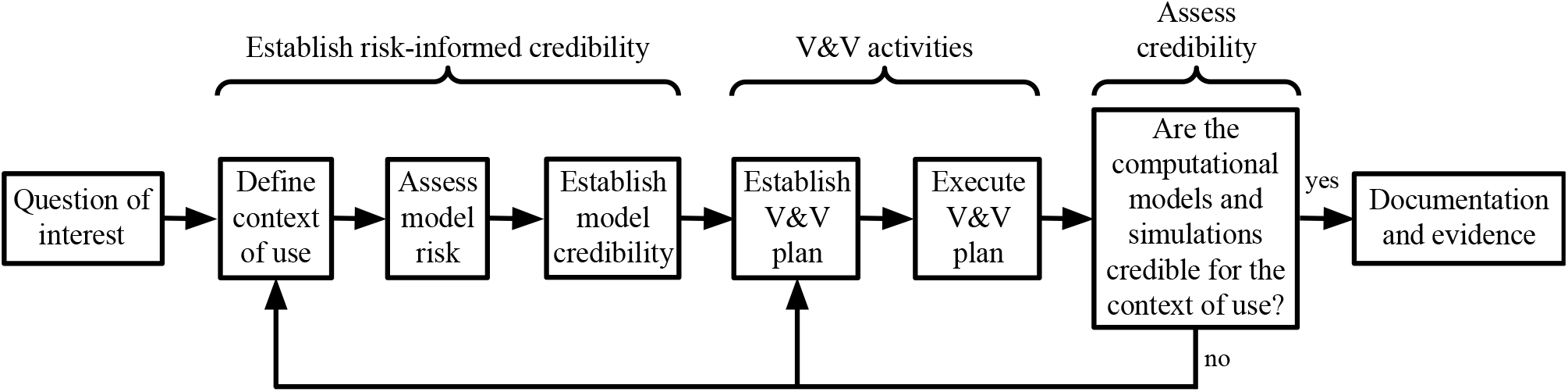
The ASME V&V40 software credibility assessment framework detailed in [34].

Considering **model risk**, the final decision consequence is high because the treatments of interest have a substantial impact on patients’ abilities (evaluated according the modified Rankin Scale [53, 54]). Model importance is estimated medium because the results are not the only factors in decision making, but they impact large artery simulation directly and functional outcome estimation indirectly in the envisaged INSIST pipeline [8]. The **verification and validation plan** relies on comparisons with analytical solutions and clinical data, respectively. Previously, preliminary validation was presented highlighting a promising agreement with clinical data regarding perfusion and infarct volume estimations during ischaemic stroke [14]. This marked the first step towards **establishing model credibility**. Beyond a thorough verification, the following sections underpin a detailed validation with multiple virtual patients by determining the parameters which ensure that the pressure, volumetric flow rate, and perfusion estimates provided by the porous FE model are accurate and physiologically realistic.

### 3.2 Spatial resolution

#### 3.2.1 Manufactured solutions

The suitability of the finite element method is evaluated based on the manufactured solution shown in Fig. 4. These synthetic solutions are reminiscent of a cortical column: the flow is driven from high pressure arterioles to low pressure venules. A uniform tetrahedral mesh is generated with two elements along each edge of the cube and uniformly refined four times to carry out a grid convergence study using first (*P*_1_) and second order (*P*_2_) elements as summarised in Tab. 1. The analytical solution and the finite element approximation along the sampling line depicted in Fig. 4 are shown in Fig. 5a, highlighting that a very good agreement is achievable. Fig. 5b presents the *L*^2^-norm defined over the entire domain based on the difference between the analytical and the numerical solutions. With increasing spatial resolution, the *L*^2^-norm decreases as expected. Superconvergence of the solution with first order elements can be observed which is a well-known feature of this FE approximation [55].

**Table 1:** Results corresponding to the grid convergence study using manufactured solutions. Normalised Grid Spacing (NGS), number of elements along each edge of the unit cube (*n*_edge_); number of elements (*n*_ele_); number of degrees of freedom (*n*_DoF_); wall time required for the iterative solution of the linear system (*t*_inv_).

| NGS | $n_{\text{edge}}$ | $n_{\text{ele}}$ | Element order | $n_{\text{DoF}}$ | $L^2$-norm | $\partial p_1 / \partial z$ at $(0.5, 0.5, 1)$ | $t_{\text{inv}}$ [s] |
| --- | --- | --- | --- | --- | --- | --- | --- |
| 16 | 2 | 48 | 1 | 81 | 0.15174 | 1.273 | 0.0042 |
| 8 | 4 | 384 | 1 | 375 | 0.08714 | 1.800 | 0.0287 |
| 4 | 8 | 3072 | 1 | 2187 | 0.02857 | 1.867 | 0.2278 |
| 2 | 16 | 24576 | 1 | 14739 | 0.00771 | 1.927 | 1.8428 |
| 1 | 32 | 196608 | 1 | 107811 | 0.00197 | 1.965 | 15.0789 |
| 16 | 2 | 48 | 2 | 375 | 0.06215 | 1.824 | 0.0161 |
| 8 | 4 | 384 | 2 | 2187 | 0.01399 | 2.006 | 0.1403 |
| 4 | 8 | 3072 | 2 | 14739 | 0.00266 | 2.008 | 1.4050 |
| 2 | 16 | 24576 | 2 | 107811 | 0.00060 | 1.999 | 16.3241 |
| 1 | 32 | 196608 | 2 | 828375 | 0.00014 | 1.998 | 162.1361 |

**Figure 4:**
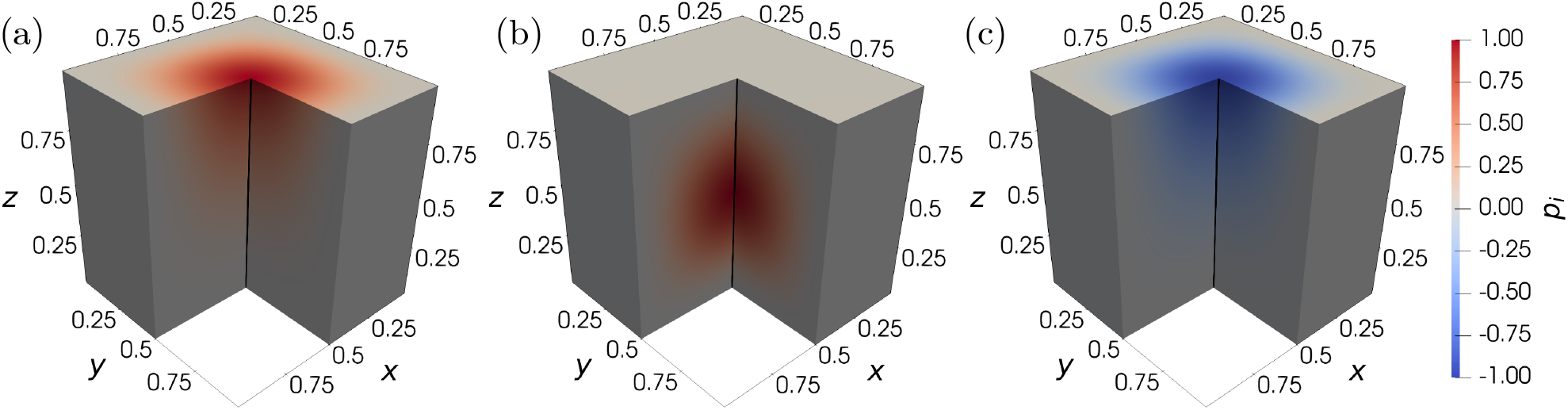
Manufactured solution in a unit cube visualised in the arteriole *i* = 1 (a), capillary *i* = 2 (b), and venule *i* = 3 (c) compartments with a sampling line (0.5, 0.5, *z*).

**Figure 5:**
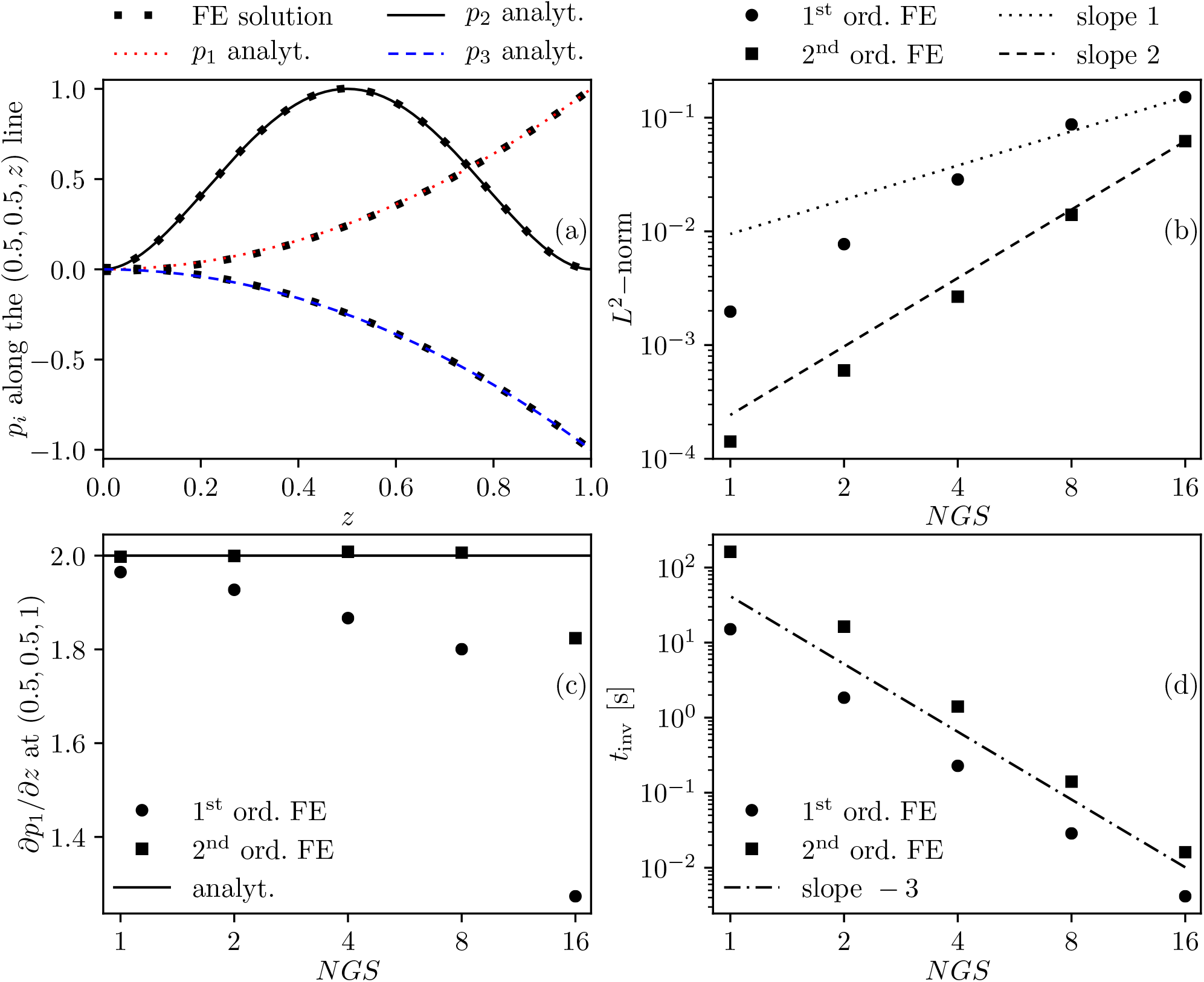
Exact manufactured solutions and FE approximations with second order (ord.) elements and normalised grid spacing (NGS) 2 along a line (a). Numerical error of the FE approximation (b), FE approximation and exact value of the arteriole gradient at a single point (c), and wall time of the linear system matrix inversion (*t*_inv_) (d) as functions of the NGS and element order.

According to Fig. 5b, solutions achieved using a second order scheme on a given mesh have about the same *L*^2^-norm as first order approximations after a single refinement, as suggested by the number of degrees of freedoms in Tab. 1. However, it is important to emphasise that the convergence of pointwise variables does not necessarily follow the convergence trend of integral quantities. Fig. 5c suggests that the second order FE scheme outperforms the first order scheme in terms of gradient estimation even if computations with the latter are carried out with a threefold refined mesh. Beyond accuracy, the computational cost of simulations as a function of the spatial resolution and the element order can be examined in Fig. 5d. In general, computations with second order elements take one order of magnitude longer and are comparable to the computational cost of simulations with first order elements on a refined mesh (see also Tab. 1).

#### 3.2.2 Whole brain model

Next, simulations are carried out with the whole brain model to investigate how uncertainty of the gradient estimation near the boundaries impacts the results. Two scenarios are considered using previously optimised settings [14] summarised in Tab. 2: baseline (healthy) and an RMCA occlusion. The algorithm used for parameter optimisation is described in Appendix B. In a porous framework, the pressure gradient is directly connected to the Darcy velocity vector which can be introduced in compartment *i* as

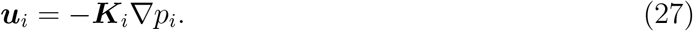

**Table 2:** List of model parameters and some reference values (mean ± standard deviation when available). The distribution of the parameters reported in [51] based on microscale vessel network simulations is not normal. The last column indicates which parameters are optimised as detailed in Appendix B.

| Parameter | Value | Reference | Unit | Optimised |
| --- | --- | --- | --- | --- |
| $k_1$ | 1.234 | 4 [22]; $0.021 \pm 0.019$ [56] | $\text{mm}^3 \text{ s kg}^{-1}$ | Yes |
| $\beta_{i,j}^G / \beta_{i,j}^W$ | 2.538 | 1.6 [22]; 1.0 [15, 16, 18] | — | Yes |
| $k_2$ | $4.28 \cdot 10^{-4}$ | $4.28 \cdot 10^{-4}$ [51]; | $\text{mm}^3 \text{ s kg}^{-1}$ | No |
| $k_3/k_1$ | 2 | 2 [22] | $\text{mm}^3 \text{ s kg}^{-1}$ | No |
| $\beta_{1,2}^G$ | $1.326 \cdot 10^{-6}$ | $1.5 \cdot 10^{-19}$ [15, 18]; $1.5 \cdot 10^{-7}$ [16];<br>$5 \cdot 10^{-6}$ [22]; $(4.13 \pm 5.49) \cdot 10^{-5}$ [56] | $\text{Pa}^{-1} \text{ s}^{-1}$ | No |
| $\beta_{2,3}^G$ | $4.641 \cdot 10^{-6}$ | $1.5 \cdot 10^{-19}$ [15, 18]; $1.5 \cdot 10^{-7}$ [16];<br>$5 \cdot 10^{-6}$ [22] | $\text{Pa}^{-1} \text{ s}^{-1}$ | No |
| CPP | 75 | $78 \pm 10$ [57] | mmHg | — |
| $\beta_{1,2}^G + \beta_{2,3}^G$ | $5.967 \cdot 10^{-6}$ | $3 \cdot 10^{-19}$ [15, 18]; $3 \cdot 10^{-7}$ [16];<br>$10^{-5}$ [22] | $\text{Pa}^{-1} \text{ s}^{-1}$ | — |

From here, the overall volumetric flow rate entering the arteriole compartment through the pial surface Γ*_P_* is simply defined as

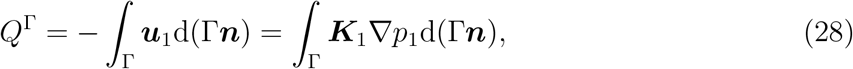

The Γ***n*** product corresponds to an outward-pointing vector with magnitude equal to the area of the Γ boundary surface. (The Darcy velocity has non-zero components perpendicularly to the boundary surface only at Γ*_P_* and therefore in Eq. (28) the integration domain Γ*_P_* is replaced with Γ for a more concise notation.)

Therefore, any uncertainty in the pressure gradient propagates directly to the volumetric flow rate. Accurate flow rate computation is important for two reasons: (i) superficial arteriole pressure and flow rates are cornerstones of a coupling scheme between the present continuum model and a network model of large arteries as detailed in Section 1; (ii) surface fluxes associated with major cerebral arteries can be directly compared with values obtained from phase-contrast MRI [58] for validation purposes.

The multi-compartment formulation offers an alternative method to compute volumetric flow rate through the brain. Introducing perfusion *F* = *β*_1,2_(*p*_1_ – *p*_2_), and integrating it over the brain volume Ω leads to

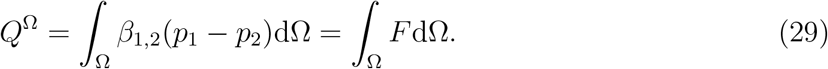

According to mass conservation *Q*^Ω^ = *Q*^Γ^ so the imbalance between the two formulations is present because the utilised FE method is not conservative. The present *H*^1^ (pressure) and *L*^2^ (velocity) spaces are selected for straightforward implementation, robustness, and relatively low memory requirements. Nevertheless, it is worth noting that numerical solutions of the Poisson equation using the mixed *H*(div) × *L*^2^ FE formulation [59, 60] are conservative, similarly to the finite volume method [61]. To mitigate the mass conservation error of the present model, two strategies are employed: (i) grid refinement; and (ii) increasing FE orders of both the pressure and the velocity. For *P*_1_ and *P*_2_ pressure elements, the natural velocity pair is *dP*_0_ and *dP*_1_ but higher order velocity approximations can be obtained by projection. The results in Tab. 3 summarise the tested cases and show that the volumetric flow rate from Eq. (29) depends on the grid size and the element order only negligibly. The error in *Q*^Ω^ should be relatively low in the investigated cases because the computed values overlap independently from the spatial resolution. For this reason, and because of the mass conservation principle, the relative difference,

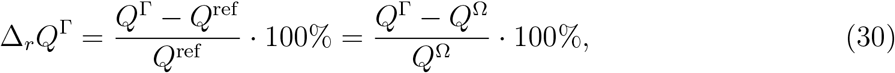

is used for error measurement.

**Table 3:** Results corresponding to the grid sensitivity analysis of the brain simulations. Due to the computational cost associated with the locally refined (loc. ref.) and the uniformly refined (unif. ref.) meshes, these simulations are run using 12 threads of an Intel Xeon E5-2640 processor and 128 GB RAM. Computations on the unif. ref. mesh with *P*_2_ elements require more than 128 GB RAM and therefore they are omitted. ^⋆^ highlights the case used to evaluate the impact of pressure and velocity inlet BCs, whereas the bold text indicates the case with the smallest difference between *Q*^Γ^ and *Q*^Ω^.

| Mesh | $n_{\text{ele}}$ | $p_i$ FE | $\mathbf{v}_i$ FE | healthy | | RMCA occl. | |
| --- | --- | --- | --- | --- | --- | --- | --- |
| | | | | $Q^\Omega$ | $Q^\Gamma$ | $Q^\Omega$ | $Q^\Gamma$ |
| baseline | 1,042,301 | $P_1$ | $dP_0$ | 604 | 464 | 493 | 380 |
| | | $P_1$ | $P_1$ | 604 | 497 | 493 | 406 |
| | | $P_2^*$ | $P_1^*$ | 600* | 559* | 478* | 447* |
| | | $P_2$ | $P_2$ | 600 | 578 | 478 | 460 |
| loc. ref. | 3,400,570 | $P_1$ | $dP_0$ | 603 | 539 | 489 | 433 |
| | | $P_1$ | $P_1$ | 603 | 546 | 489 | 439 |
| | | $P_2$ | $P_1$ | 600 | 582 | 476 | 461 |
|  |  | **$P_2$** | **$P_2$** | **600** | **597** | **476** | **471** |
| unif. ref. | 8,338,408 | $P_1$ | $dP_0$ | 602 | 511 | 486 | 411 |
| | | $P_1$ | $P_1$ | 602 | 527 | 486 | 425 |

Figs. 6a and b show the imbalance between *Q*^Ω^ and *Q*^Γ^ highlighting that without improved ***u**_i_* approximation the surface flux is always underestimated by *Q*^Γ^. Both refinement and increased FE order can mitigate the error of *Q*^Γ^. However, higher order pressure and velocity approximations are required to keep |Δ_*r*_*Q*^Γ^| < 5%. Fig. 6c displays the wall time of the simulations emphasising the increased computational cost required for more accurate flux estimations. In summary, second order velocity and pressure elements on a relatively coarse mesh (about 10^6^ elements) can provide a reasonable compromise between accuracy and performance if computing *Q*^Γ^ is important. Otherwise, perfusion and pressure distributions can be estimated reasonably well on coarse meshes with about 1 million elements.

**Figure 6:**
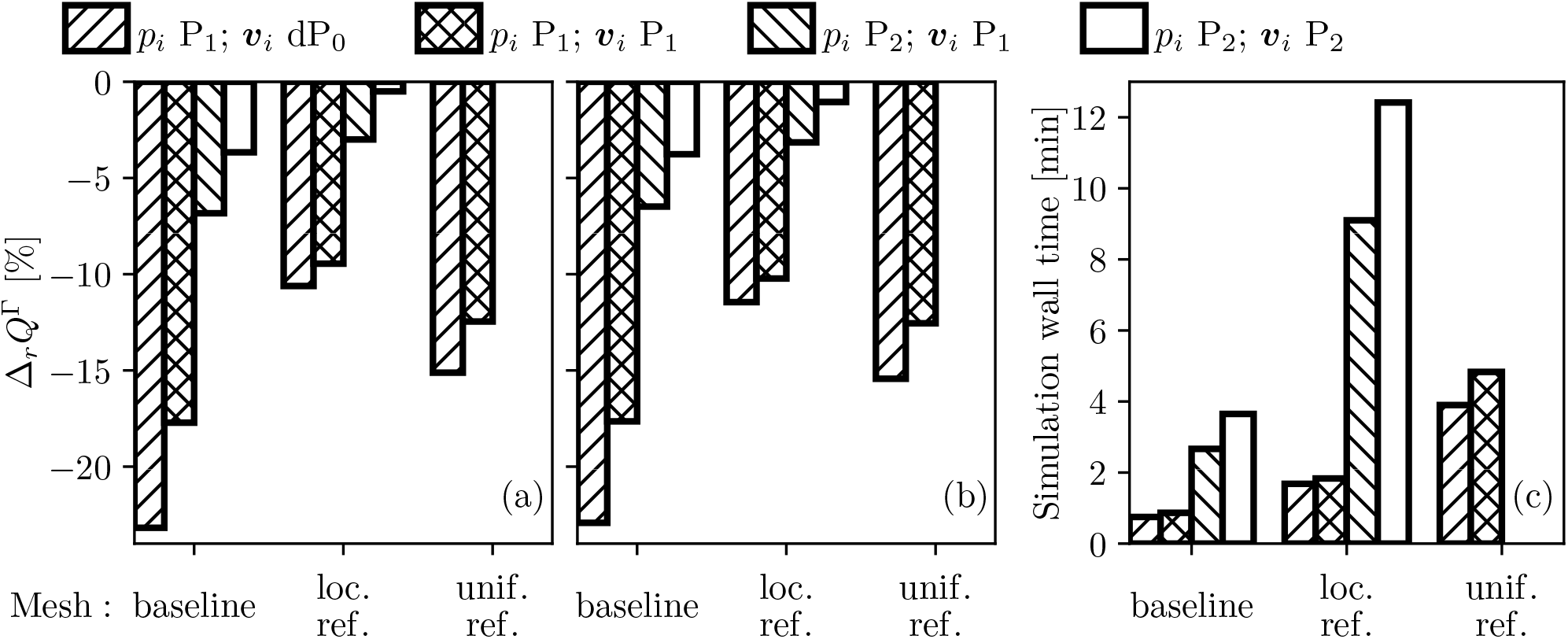
Relative error of the superficial volumetric flow rate estimation as functions of the spatial resolution and the FE order in the case of the baseline (a) and RMCA occlusion (b) scenarios. Computational time averaged between the healthy and the occluded cases (c). The “loc. ref.” and “unif. ref.” abbreviations correspond to locally and uniformly refined meshes, respectively (see Tab. 3 for further details).

Once it is established under what conditions the present 3D model can provide accurate estimation of volumetric flow rate through cortical surface regions, it is possible to compute volumetric flow rate of the major cerebral arteries. Simulation results are compared with values from a clinical study in Tab. 4. Whereas the model is optimised for overall brain perfusion, blood flow rate through these major arteries provides a chance for independent validation. It is promising that volumetric flow rate in both the ACA and the MCA is predicted within one standard deviation of the experimental values. Volumetric flow rate in the PCA is somewhat overestimated. This simple comparison is an important milestone in the quantitative validation of the 3D brain model and provides guidance about development directions for the future.

**Table 4:** Comparison of some model parameters and results with literature data: volumetric flow rates of the anterior (ACA), middle (MCA) and posterior (PCA) cerebral arteries obtained by surface integration of the velocity vector over perfusion territories; cortical surface area (*A*_cortical_) and brain volume (*V_b_*).

|  | Simulation | Reference |
| --- | --- | --- |
| $Q$ ACA/MCA/PCA [ml/min] | 66 / 126 / 67 | $75 \pm 15$ / $131 \pm 23$ / $51 \pm 10$ [58] |
| $Q$ total [ml/min] | 600 | $657 \pm 94$ [58]; $484 \pm 88$ [62] |
| $A_{\text{cortical}}$ [cm<sup>2</sup>] | 1005 | 1770 [63] |
| $V_b$ [ml] | 1390 | $869 \pm 99$ [64] |

The results emphasise the importance of accurate mapping between voxelised vascular atlases and the triangulated brain surface and draw attention to geometrical flaws. The brain model does not capture the lateral sulcus and the insula. Because of these neglected regions, the brain surface is underestimated whereas the brain volume is overestimated (see Tab. 4). It is anticipated that the 3D porous model will provide more reliable volumetric flow rate estimations once these geometrical issues are overcome. However, obtaining accurate discretised brain geometries suitable for FE computations remains a challenging task, partially because of labelling the boundary regions, and therefore such efforts are left to a future study.

### 3.3 Boundary conditions

Results from the previous section draw attention to issues related to flux computation in a porous FE framework. Instead of constant pressure, uniform velocity has been used as an inlet BC [15–19], where the velocity is calculated as the ratio of a given volumetric flow rate and the inlet surface area. This NBC at the inlet might distort the results because of errors related to gradient estimation so we next investigate the difference between pressure and velocity inlets using the present model. To this end, a single simulation with constant inlet velocity is run with the same settings as the case in Tab. 3 marked with ^⋆^ so that the total inlet volumetric flow rate remains unchanged.

*Q*^Ω^ = 600 [ml/min] is preserved, indicating that the simulations with pressure and velocity inlets are equivalent regarding average brain perfusion. *Q*^Γ^ is impacted only slightly: pressure inlet leads to 559 [ml/min] whereas velocity inlet results in 572 [ml/min] (both values should be 600 [ml/min] based on mass conservation). The effects of the inlet boundary condition on other statistics regarding the pressure and the perfusion fields are shown in Fig. 7. Most of the volume averaged pressure (〈*p_i_*〉) and perfusion (〈*p_i_*〉) values are altered only slightly by the inlet boundary conditions. However, extrema of both the pressure and the perfusion fields change substantially when the inlet velocity BC is used. Minimum values are underestimated so that the relative difference of the minimum arteriole pressure and perfusion values are approximately – 100% because min *p*_1_ and min *F*) are close to zero when uniform inlet velocity BCs are used. By comparison, the inlet velocity BCs lead to overshot maximum values nearly by a factor of 10 compared to the case with inlet pressure BCs (see max *p*_1_ and max *F* in Fig. 7).

**Figure 7:**
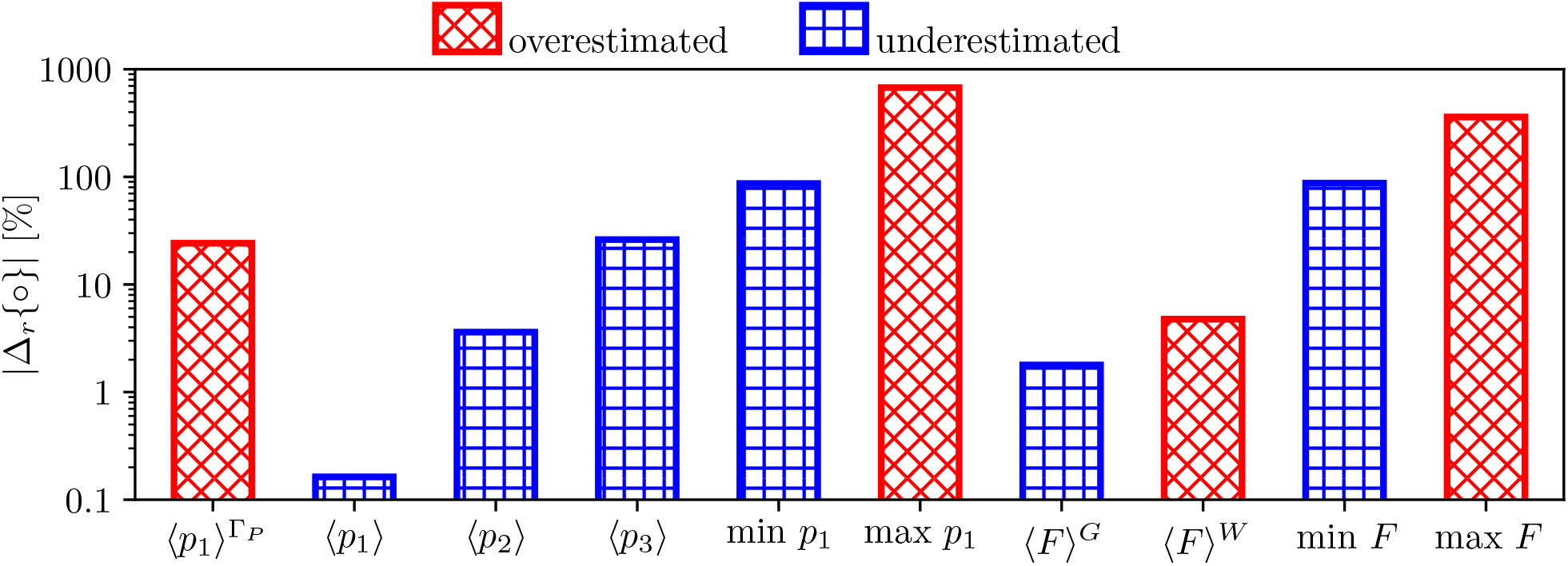
Absolute value of the relative difference between some statistics extracted from simulations carried out with inlet pressure and velocity BCs. Average arteriole pressure over the pial surface 〈*p*_1_〉^Γ*_P_*^; average pressure in the arteriole (〈*p*_1_〉), capillary (〈*p*_2_〉), and venule (〈*p*_3_〉) compartments over the entire brain; minimum (min) and maximum (max) arteriole pressure (*p*_1_) and perfusion (*F*); average perfusion in grey (〈*F*〉*^G^*) and white (〈*F*〉*^W^*) matter. The colour and pattern of each bar indicates whether results with the inlet velocity BCs overshoot or underestimate the reference values corresponding to the pressure inlet BCs.

In Figs. 8a-d, and b-e, the pressure and perfusion fields corresponding to inlet pressure and velocity BCs, respectively, emphasise further the difference between the two solutions. The arteriole velocity magnitude displayed over the cortical surface (Fig. 8c) shows that in the case of pressure inlet BC the velocity is strongly inhomogeneous. It is a remarkable feature of this scenario that high velocity values are predicted along the major cerebral arteries branching from the circle of Willis because the whole brain model does not include any information about the location of these vessels. The solution obtained with constant pressure BC suggests that descending arterioles branching from the major cerebral arteries play a key role in providing well-balanced grey and white matter perfusion. The velocity distribution over the cortical surface is not known *a priori* and therefore it is challenging to impose feasible velocity distributions with velocity inlet BC.

**Figure 8:**
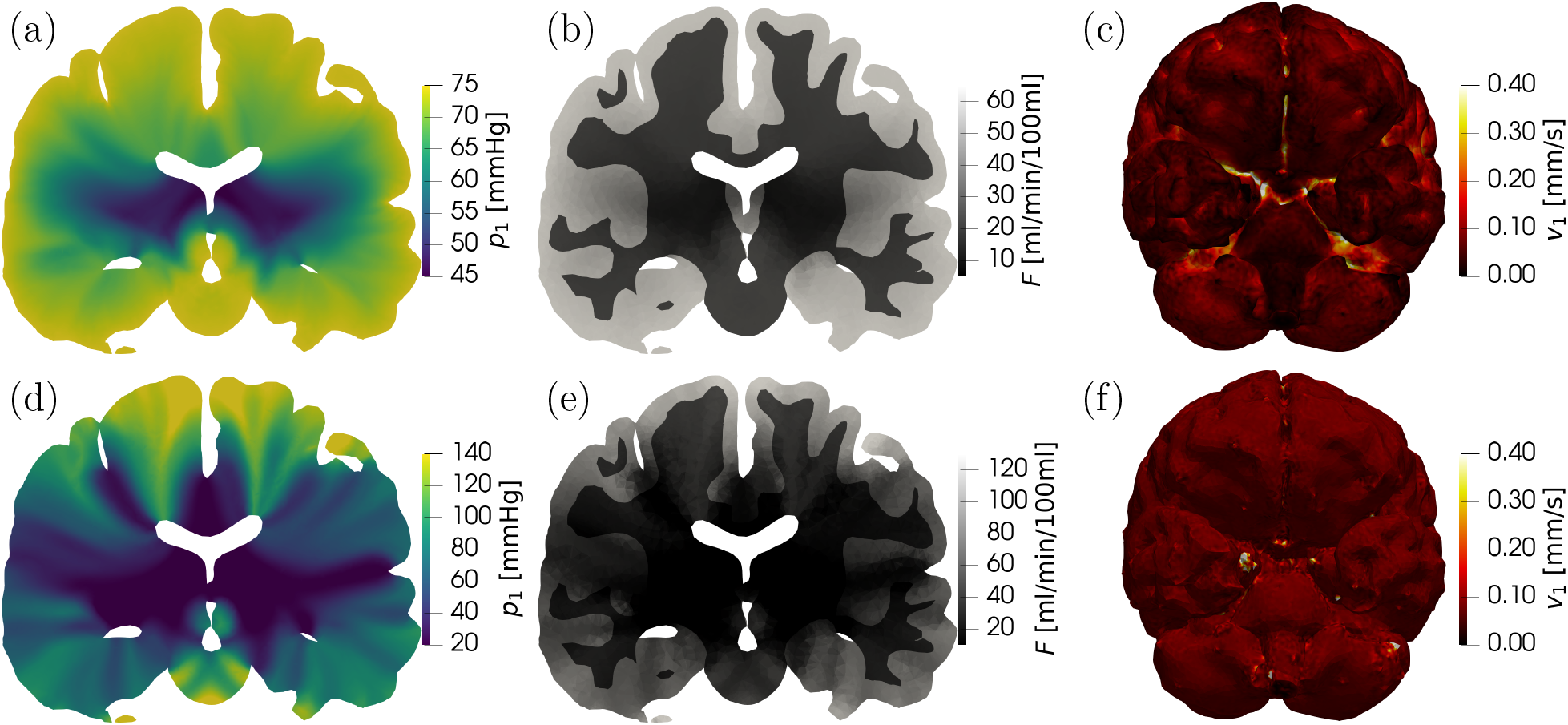
Pressure *p_a_* (a-d) and perfusion *F* (b-e) distributions along a coronal plane, and velocity magnitude (c-f) along the cortical surface. Constant pressure inlet (a-c) and constant velocity inlet (d-f). In (f), the non-uniform velocity magnitude is associated with numerical errors. Results correspond to the mesh with local grid refinement at the boundaries and *P*_2_ pressure and velocity elements (details in Tab. 3).

The volumetric flow rates corresponding to boundary regions of the major cerebral arteries are summarised in Tab. 4. Based on previous clinical investigations [58, 62], it is clear that each major cerebral artery contributes differently to volumetric blood flow rate to the brain. Even if we assume that superficial perfusion territories of cerebral arteries are proportional to the associated blood flow rate, it is important to recognise that the volumetric flow rate should be heterogeneous as dictated by underlying grey and white matter volumes. Although the validity of homogeneous inlet velocity is physiologically questionable, prescribing constant pressure inlets in healthy scenarios is supported indirectly by clinical data. Based on ASL MRI [47], cerebral perfusion regions have been found to be well-separated indicating that the pressure gradient between these territories is small. Furthermore, dynamic computed tomography angiograms indicate that flow through leptomeningeal collaterals is activated as a result of cerebral artery occlusion [65]. We hypothesise that in healthy cases, leptomeningeal collaterals [2, 50, 65–68] as well as pial arterioles [69–71] act to equalise blood pressure in descending arterioles near the cortical surface. This hypothesis could be tested once the present perfusion model and the 1D model incorporating the larger arteries [13] are coupled, and it is supported by pressure data obtained in rodents which suggest small pressure drop in large arteries compared to the microcirculation [72–74].

In summary, there is no evidence that large arteries distribute blood flow uniformly along the cortical surface. Prescribing uniform inlet velocity as a boundary condition in the arteriole compartment is questionable, and leads to extreme pressure and perfusion values out of the expected range. Based on anatomical and physiological considerations, uniform inlet pressure appears to be a more suitable boundary condition in healthy scenarios.

### 3.4 Sensitivity analysis

Determining the actual values of parameters in physiological models is a challenging task because of limited experimental data, high patient-specific variability, and the typically high dimensionality of the parameter spaces. Considering porous cerebral haemodynamics, this issue is far from being settled as indicated by the broad range of reported parameters summarised in Tab. 2. With increasing computational costs, mapping a high dimensional parameter space quickly becomes unfeasible. Beyond parameter optimisation, sensitivity analysis and the associated uncertainty quantification are other tasks which require large batches of simulations. Here we aim to utilise the one-dimensional model described in Section 2.3 in order to ease these tasks.

#### 3.4.1 One-dimensional brain tissue column

The one-dimensional multi-compartment porous model enables the investigation of tissue columns including both grey and white matters so that the corresponding model parameters are identical with the ones used for the three-dimensional brain model (Tab. 2). In the 1D case, *x* = 0 and *x* = *l* correspond to the pial and the ventricular surfaces, respectively. Therefore, the same boundary conditions can be imposed in the 1D tissue column and the 3D brain. (Similarly to the 3D brain, pressure values in the 1D case are presented relative to the venous outlet pressure *p_v_*.) Thereafter, the one-dimensional model has only two free parameters: the lengths of the grey (*l_G_*) and the white (*l_W_*) matter domains defined so that *l* = *l_G_* + *l_W_*. The equivalent *l_G_* = 13.55 and *l_W_* = 7.99 [mm] is uniquely determined by the human brain model used here and can be obtained by optimisation. The corresponding cost function is zero if the one-dimensional model returns 〈*F*〉 = 〈*F*〉*^G^* = 56, and 〈*F*〉*^W^* = 21 [ml/min/100ml] which overlap exactly with volume-averaged perfusion values of the three-dimensional brain model.

Fig. 9 displays the analytical pressure distributions in the arteriole, capillary and venule compartments. A one-dimensional FE approximation with 100 *P*_1_ elements is also shown which is in excellent agreement with the analytical results. The average pressure values provided by the 1D model are within ±4% of those computed using the 3D brain in every case. One-dimensional simulations are approximately 3000 times faster compared to their 3D counterparts.

**Figure 9:**
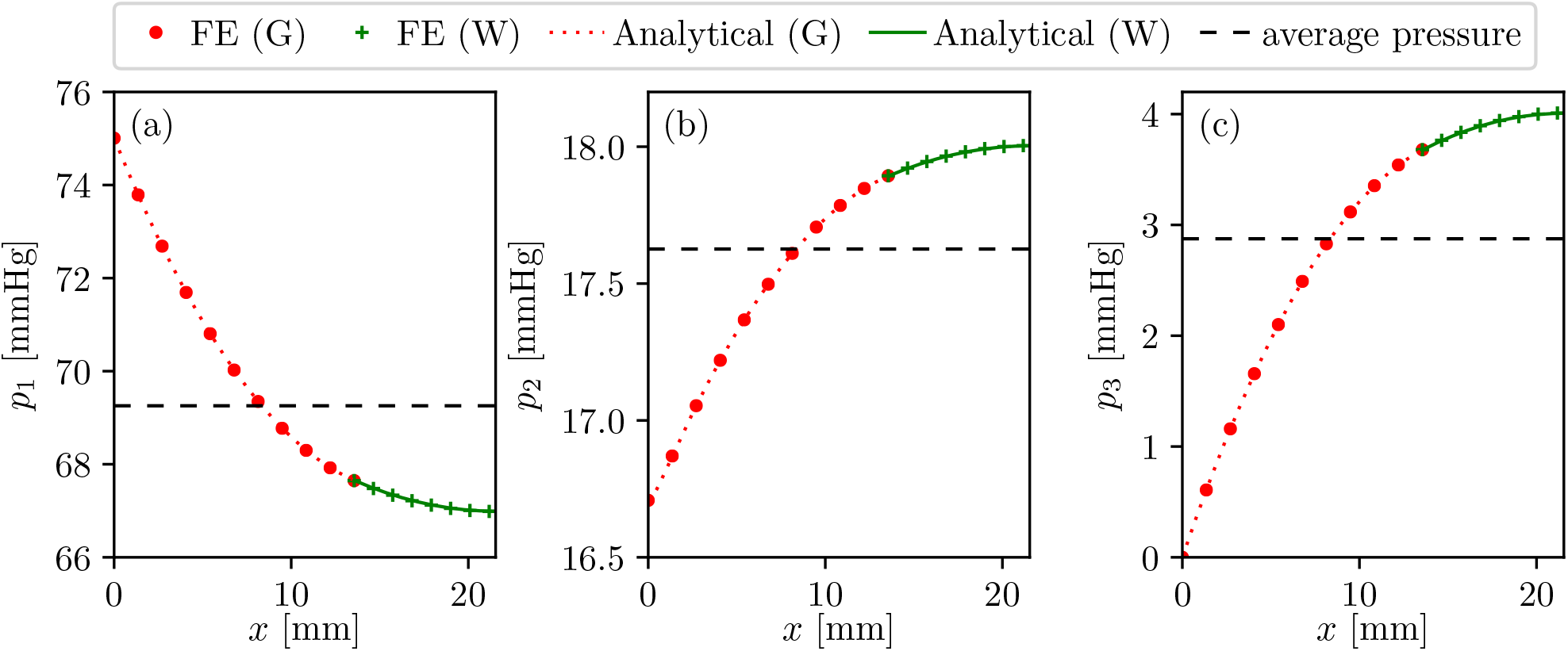
Analytical and numerical solutions of the one-dimensional problem representing a brain tissue column perpendicular to the cortical surface: arteriole (a), capillary (b) and venule (c) compartments. Results in both grey (G) and white (W) matter are displayed.

#### 3.4.2 Mapping the parameter space

The 1D model is a good candidate for parameter space mapping because of its low computational cost. A disadvantage of the analytical formulation is that with the parameters presented in Tab. 2, the linear equation system governing the boundary conditions is very stiff, probably because the parameters have different orders of magnitude. The condition number depends nonlinearly on the reference length scale used for the nondimensionalisation. To avoid the issue of suitable reference length scale selection, we will use the 1D finite element model verified in Fig. 9 for a detailed sensitivity analysis.

Next, each parameter is perturbed independently from others. The one-at-a-time approach is applicable and the corresponding sensitivity analysis is representative of the system behaviour because the governing equation set (1) is linear [33]. The parameter range is [10%;1000%] compared to the values in Tab. 2 with 101 samples along each dimension. In addition, a geometrical scaling factor is introduced based on the ratio of the original and the perturbed volume of the computational domain. In order to evaluate whether the 1D model can capture the system behaviour, 11 samples along each dimension of the parameter space are tested with the 3D brain model using *P*_1_ pressure elements.

Results are summarised in Fig. 10 showing brain perfusion as a function of the model parameters in the healthy scenario. Based on the 3D simulations, RMCA occlusion is also considered so that the perfusion in the occluded scenario and infarcted volume change are also displayed. Infarcted volume is estimated solely based on the perfusion distribution. A region with more than 70% perfusion drop compared to the baseline scenario is identified as an infarcted zone inspired by perfusion imaging-based methods [75, 76]. The obtained value estimating infarcted volume is used here solely as an indicator of perfusion drop severity because this method overestimates the ischaemic region compared to diffusion-weighted magnetic resonance imaging [77].

**Figure 10:**
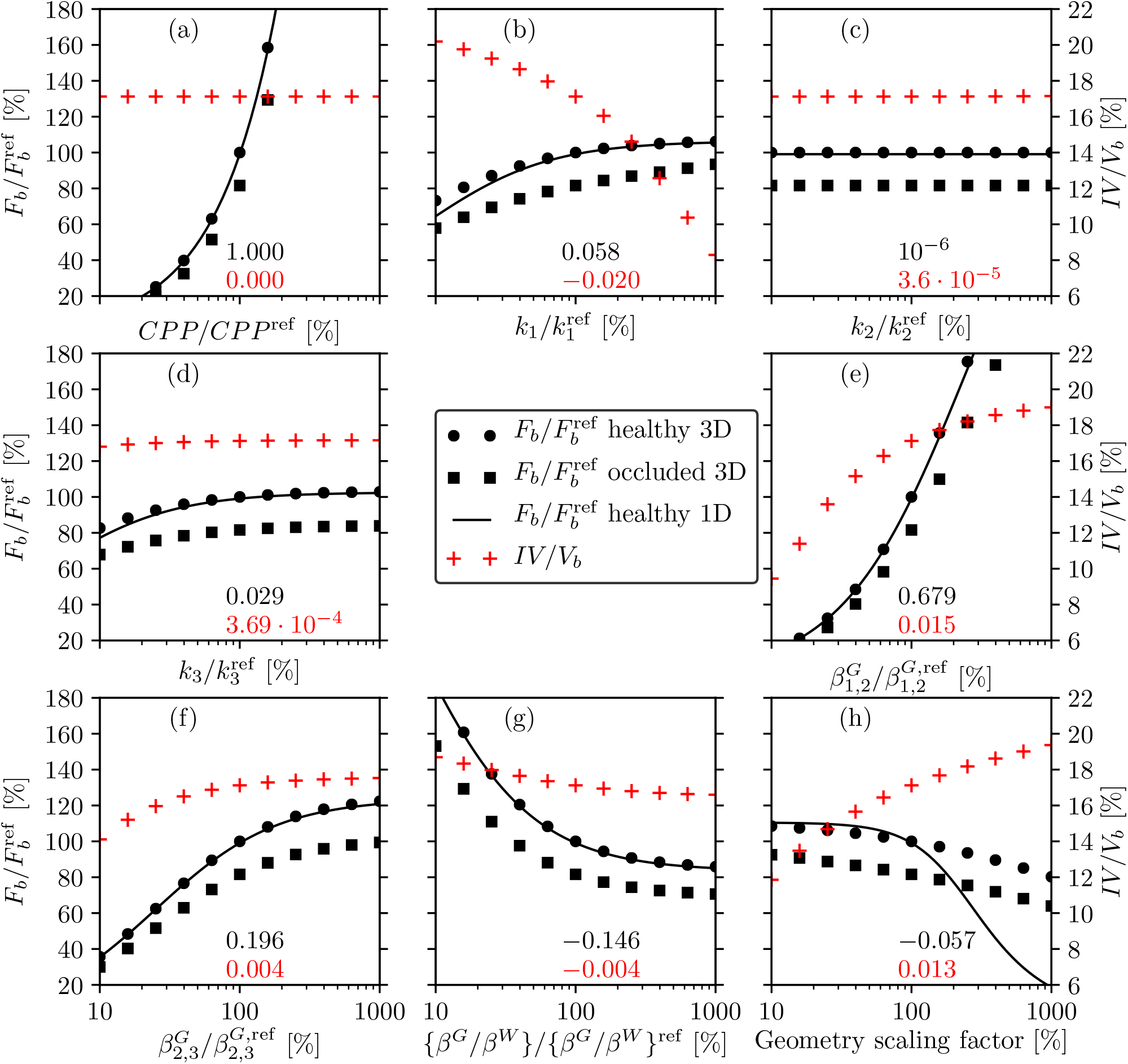
Sensitivity analyses of the 1D and 3D models. Brain perfusion (left axes) and infarcted volume fraction (right axes) as functions of change in the following parameters: cerebral perfusion pressure (a); arteriole (b), capillary (c), and venule (d) permeabilities; arteriole-capillary (e) and capillary-venule (f) coupling coefficients in the grey matter; ratio of grey and white matter coupling coefficients (g); geometry scaling factor (h). The top (black) and bottom (red) numbers on each subplot stand for the sensitivity of healthy brain perfusion and infarcted volume fraction, respectively. Reference values of the parameters are listed in Tab. 2.

According to Figs. 10a-h, the 1D solutions follow the trend of the 3D simulations in a wide parameter range in every case except the geometrical scaling factor. Considering that the present 1D model does not account for curvature effects, the results could potentially be improved with a cylindrical or spherical formulation. The sensitivity of each parameter is estimated as the normalised partial derivative of brain perfusion and infarcted volume at 100% of the abscissa (around baseline values presented in Tab. 2). The derivatives computed based on the 3D simulations in Fig. 10a-h are listed above the abscissa and indicate the percentage change of brain perfusion and infarcted volume as a response to unit percentage change in model parameters.

In terms of haemodynamics, it should be noted that the present model captures solely the resistive part of the microcirculation and neglects conductance and inductance associated with vessel wall stiffness and pulsatility. The linear relationship between brain perfusion and perfusion pressure is a direct manifestation of the “hydraulic Ohm’s law” (Fig. 10a). As other parameters are increased, brain perfusion tends to exhibit a saturating behaviour even if it is not obvious within the displayed parameter range (Fig. 10b-g) Considering the infarcted volume fraction, the insensitivity to CPP change originates from its definition restricted to relative perfusion change (as CPP is increased perfusion in the healthy and the occluded cases increases by the same amount).

Perturbations in the capillary and venous permeabilities have a relatively weak impact on both *F_b_* and *IV*. Fluid flow within the capillary compartment is relatively small. This result can be interpreted as flow “avoiding” the capillaries because of their high resistance, which overlaps with detailed network simulation of the rat microcirculation [74]. Considering an indefinite domain with solely grey or white matter and constant but different arteriole and venous pressures, blood flow is non-zero between the compartments but it is zero inside each compartment. In the microcirculation, capillaries cannot be bypassed so it is important to recognise that the coupling coefficients must account not only for pre- and post-capillaries but also for those capillary vessels which establish the shortest route between arterioles and venules. Therefore, the results show strong sensitivity to changes in coupling coefficients, which are responsible for the majority of the system resistance and consequently most of the pressure drop.

The geometry scaling factor has a strong influence on the infarcted volume fraction. If one of the major feeding arteries is occluded, it seems logical that for a given purely resistive vasculature a larger brain suffers more compared to a smaller brain. Such effects might be observable in clinical data but they are unlikely to be statistically significant compared to other factors, such as the extent of collateralisation. Nevertheless, the results suggest that a representative brain geometry is essential to keep the numerical uncertainty of the simulations low. In addition, the data presented in Fig. 10, and the one-dimensional model particularly, can help to accelerate virtual patient generation, where parameter sets must be found so that the resulting perfusion distribution is representative of a patient cohort.

The sensitivity analysis shown in Fig. 10 emphasises the importance of the (i) boundary conditions and (ii) coupling coefficients. Based on the literature data in Tab. 2, the coupling coefficients are burdened by the highest uncertainty which can undermine the applicability of the present model in terms of perfusion and infarct volume estimation through uncertainty propagation.

### 3.5 Limitations

This subsection summarises some of the most important overlooked factors in the present study. Investigations are limited thus far to a single patient-specific brain geometry because of challenges associated with mesh generation. The porous model relies on multiple scale separation even though the vessel diameter in the vasculature changes continuously [20]. The present study demonstrated promising quantitative validation based on blood flow rate in major cerebral arteries but a more comprehensive validation is essential to ensure wider, multi-purpose applicability of the model.

The study is restricted to a porous model with three compartments (arteriole, capillary, venule) which are connected by homogeneous coupling coefficients both in grey and white matter. The model cannot represent arterioles which branch relatively far away from the cortical surface and hence feed deeper tissue regions. For this reason, low perfusion regions caused by occluded cerebral arteries are always connected to the brain surface so that isolated white matter infarction near the ventricles cannot be captured. On the one hand, such effects could be resolved by model expansion, for instance, by introducing two arteriole compartments which perfuse grey and white matter separately and rely on different boundary conditions. On the other hand, models with a single or two [22] compartments might be found more suitable in certain cases. Considering that the present study focuses on the sensitivity analysis of a three-compartment brain model for acute ischaemic stroke, exploring alternative porous frameworks for other specific (patho)physiological problems is left to future investigations.

From the haemodynamics point of view, the present model is purely resistive and aims to capture statistically steady state (time-averaged) brain perfusion. Therefore, the following physiological phenomena have been neglected: vessel wall stiffness (conductance), unsteady flow phenomena (inductance) [78], and cerebral autoregulation [79]. The associated processes and mechanisms are strongly time-dependent and often rely on nonlinear processes leading to significant complexity which is beyond the limits of the present study. Nevertheless, the authors hope that the present work will contribute to the creation of a comprehensive cerebral blood flow model which can incorporate these effects, in addition to pathophysiological processes, such as cerebral oedema [30], emboli advection and blockage of the microcirculation [56, 80, 81], and spreading of ischaemic tissue damage [52, 82, 83].

## 4 Conclusions

The present study set out to verify and explore the sensitivity of a porous brain perfusion model and to prepare the ground for validation and uncertainty quantification as specified in the ASME V&V40 standard [34]. To this end, simulations were carried out using the finite element method and three test cases: a unit cube with manufactured solutions, a human brain geometry, and a one-dimensional brain tissue column. For the unit cube and the brain tissue column, analytical solutions were presented and used for verification. This approach allowed direct comparison between analytical and numerical results and enabled the identification of certain settings which were found crucial to minimise numerical errors.

Based on the key findings of this study, the following guidelines are established regarding the efficient usage of three-dimensional porous finite element models for steady state cerebral perfusion simulation:

- Blood pressure and perfusion can be estimated reasonably well on a relatively coarse grid and first order pressure elements.
- In order to approximate volumetric flow rate through superficial territories accurately with minimal computational effort, higher order finite elements are essential. Volumetric flow rate estimation can be further improved with local grid refinement in the vicinity of the superficial territories.
- Porous models can provide a good estimation of blood flow rate through major cerebral arteries when higher order elements and constant inlet pressure boundary conditions are used.
- Using uniform velocity inlet boundary conditions does not have solid physiological foundations because blood flow is likely not distributed proportionally to the cortical surface area. However, when considering healthy scenarios, constant pressure inlets can be prescribed with higher confidence.
- A multi-compartmental porous system can be solved analytically in one-dimensional cases. After determining the equivalent thickness of the subdomains, the one-dimensional system behaviour is representative of the three-dimensional organ-scale model. The one-dimensional case is suitable for parameter mapping and sensitivity analysis at a computational cost three orders of magnitude smaller compared to three-dimensional models.

The results lay down the foundations for validation, uncertainty quantification, and for the development of a multiscale model which incorporates large arteries as well as the microcirculation. The present work contributes to the creation of a reliable computational model which can assist in the design of clinical trials and in clinical decision making related to cerebrovascular diseases.

## Acknowledgements

This work was funded by the European Union’s Horizon 2020 research and innovation programme, the INSIST project, under grant agreement No 777072. Thanks are due to the members of the INSIST consortium for useful discussions and their sustained support.

## Authors’ contributions

TIJ carried out the research and prepared a draft of the manuscript under the supervision of SJP. RMP developed the brain territory clustering algorithm under the supervision of AGH. TIJ, WKE-B, and SJP designed the research and every author contributed to the revision of the paper.

## Conflict of interest declaration

The authors declare that they have no conflict of interest.

## Appendix A Permeabilities and coupling coefficients for the manufactured solutions

To obtain a synthetic pressure field, the permeability tensors are set to

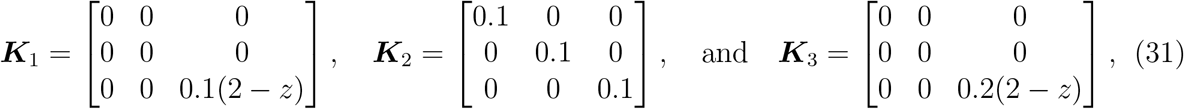

whereas the non-zero elements of the coupling coefficient matrix are

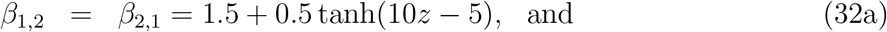

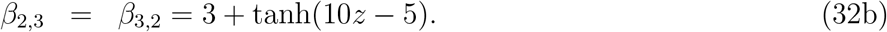

The hyperbolic tangent functions in the elements of the coupling coefficient matrix are selected to capture the sudden change in the material properties of grey and white matter. ***K**_i_* and *β_i,j_* are approximated by fourth order polynomials for the finite element computations.

## Appendix B Whole brain model parametrisation

The baseline model parameters listed in Tab. 2 are determined using the method established in [14] as summarised below.

1. The capillary permeability *k*_2_ and the venule-arteriole permeability ratio *k*_3_/*k*_1_ are assumed to be given based on previous studies [22, 51];
2. The ratio of the grey matter coupling coefficients (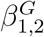 and 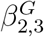) can be calculated based on the spatially averaged pressure values in the grey matter (〈*p_i_*〉*^G^*) using [14]

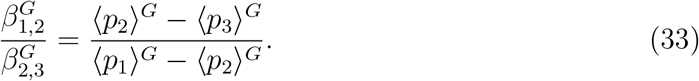 The ratio of the pressure drops is inferred from rodent experiments [72, 73]. Thereafter, both 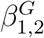 and 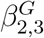 can be calculated once the cerebral perfusion pressure is estimated [57] and grey matter perfusion 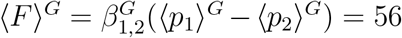 [ml/min/(100 ml)] is selected.
3. The remaining two parameters, namely *k*_1_ and 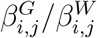, are obtained using optimisation [84, 85] relying on the minimalisation of the

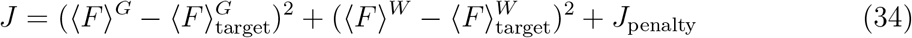

cost function. The target perfusions in grey and white matter are set to physiologically realistic values [86]: 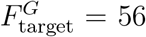 and 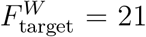 [(ml blood)/min/(100 ml tissue)]. Furthermore, a penalty term (*J*_penalty_) is applied to restrict the minimum and maximum perfusion values:

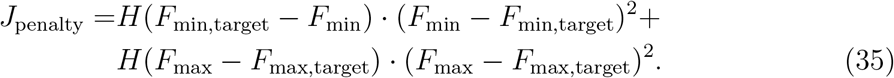 Here, *H* is the Heaviside function resulting in a non-zero *J*_penalty_ only if the extrema are out of the *F*_min,target_ = 10 and the *F*_max,target_ = 80 [(ml blood)/min/(100 ml tissue)] range.

## Footnotes

1 Einstein summation convention is not utilised in this study.

## Notes

### Competing Interest Statement

The authors have declared no competing interest.

### Summary of Updates

Figure 1 has been added to explain the physiological principles regarding the porous microcirculation model. Figure 2 has been updated to avoid overlap with previously published materials. Figure 3 and Section 3.1 have been added to reflect on the V&V40 standard of ASME.

